# Exceptional Solvent Tolerance in *Yarrowia lipolytica* Is Enhanced by Sterols

**DOI:** 10.1101/324681

**Authors:** Caleb Walker, Seunghyun Ryu, Cong T. Trinh

## Abstract

Microbial biocatalysis in organic solvents such as ionic liquids (ILs) is attractive for making fuels and chemicals from complex substrates including lignocellulosic biomass. However, low IL concentrations of 0.5-1.0 % (v/v) can drastically inhibit microbial activity. In this study, we engineered an exceptionally robust oleaginous yeast *Yarrowia lipolytica*, YlCW001, by adaptive laboratory evolution (ALE). The mutant YlWC001 shows robust growth in up to 18% (v/v) 1-ethyl-3-methylimidazolium acetate ([EMIM][OAc]), which makes it the most IL-tolerant microorganism published to our knowledge. Remarkably, YlCW001 exhibits broad tolerance in most commonly used hydrophilic ILs beyond [EMIM][OAc]. Scanning electron microscopy revealed that ILs significantly damage cell wall and/or membrane of wildtype *Y. lipolytica* with observed cavities, dents, and wrinkles while YlCW001 maintains healthy morphology even in high concentrations of ILs up to 18% (v/v). By performing comprehensive metabolomics, lipidomics, and transcriptomics to elucidate this unique phenotype, we discovered that both wildtype *Y. lipolytica* and YlCW001 reconfigured membrane composition (e.g., glycerophospholipids and sterols) and cell wall structure (e.g., chitin) under IL-stressful environments. By probing the steroid pathway at transcriptomic, enzymatic, and metabolic levels, we validated that sterols (i.e., ergosterol) are a key component of the cell membrane that enables *Y. lipolytica* to resist IL-responsive membrane damage and hence tolerate high IL concentrations. This study provides a better understanding of exceptional robustness of *Y. lipolytica* that can be potentially harnessed as a microbial manufacturing platform for production of fuels and chemicals in organic solvents.

## Significance

Robustness is an important production phenotype for any industrial microbial catalyst to acquire but it is complex and difficult to engineer. Through adaptive laboratory evolution, in combination with comprehensive omics analysis, we shed light on the underlying mechanism of how *Y. lipolytica* restructures its membrane to tolerate high levels (up to 18% v/v) of ILs, which makes our evolved strain the most IL-tolerant microorganism reported to date. Specifically, we discovered that sterols play a key role for enhancing exceptional IL tolerance in *Y. lipolytica*. Overall, this study provides fundamental understanding and engineering strategy for exceptional robustness of oleaginous yeasts to improve growth and novel biotransformation in inhibitory organic solvents.

## Introduction

Robustness is an important phenotype for any industrial microbe to acquire for biosynthesis of desirable molecules (1–3). Recently, microbial biocatalysis in green organic solvents such as ionic liquids (ILs) has become attractive to produce biofuels (4, 5) and high-value bioproducts (6–8). One key advantage is that ILs can effectively process complex and recalcitrant substrates such as lignocellulosic biomass for microbial fermentation (9–13). Further, these ILs can function as extractants for *in situ* separation of desirable molecules (14–18). Thus, it is highly desirable to harness novel microbes for biocatalysis in organic solvents.

To perform novel biotransformation in ILs, microbes need to be robust to tolerate high concentrations of solvents (19, 20). These solvents are inhibitory because they severely disrupt cell membranes and intracellular processes (21–23). For applications such as simultaneous saccharification and fermentation of IL-pretreated lignocellulosic biomass, microbes that are active in high IL-containing media are desirable because expensive IL washing and recycling steps prior to fermentation are not needed (7, 24). Unfortunately, most industrially-relevant platform organisms, *Escherichia coli* and *Saccharomyces cerevisiae*, are severely inhibited in IL-containing media even at low concentrations, for instance, 1% (v/v) [EMIM][OAc] (25, 26).

To overcome IL toxicity, various targeted and evolutionary engineering strategies have been performed. For instance, an *E. coli* strain was engineered to produce desirable chemicals from IL-pretreated biomass in the presence of 100 mM (< 2% v/v) [EMIM][OAc] by mutating the endogenous transcriptional regulator RcdA (P7Q) to de-repress expression of the inner membrane pump YbjJ (27) or overexpressing a heterologous IL-specific efflux pump, EliA, isolated from *Enterobacter lignolyticus* (28, 29). Likewise, adaptive laboratory evolution (ALE) was conducted to generate an *E. coli* mutant with robust growth in 500 mM (8.3 % (v/v)) [EMIM][OAc] (30). By deleting a highly IL-responsive, mitochondrial serine/threonine kinase gene *ptk2* (which activates a plasma-membrane proton efflux pump Pma1), an engineered *S. cerevisiae* strain became capable of tolerating > 2% (v/v) [EMIM][Cl] (31). Even though the above engineering strategies resulted in enhanced IL-tolerant phenotypes, the engineered strains are not tolerant enough to robustly grow in in 10 % (v/v) ILs unlike *Y. lipolytica.* This highlights the complexity of IL-tolerant phenotypes and the limited understanding of IL-tolerant mechanisms.

Synergistically, exploring microbial and genetic diversity can potentially discover novel genotypes conferring novel IL tolerance that typically does not exist in the current platform organisms. Bioprospecting the tropical rainforest soil resulted in isolation of a novel *Enterobacter lignolyticus* that could survive in 0.5 M (6.6 % (v/v)) [EMIM][Cl] (29). Likewise, screening microbial diversity from a collection of 168 fungal yeasts identified 13 robust strains that can tolerate up to 5% (v/v) [EMIM][OAc], including *Clavispora*, *Debaryomyces*, *Galactomyces*, *Hyphopichia*, *Kazachstania*, *Meyerozyma*, *Naumovozyma*, *Wickerhamomyces*, *Yarrowia*, and *Zygoascus* genera (20). Among the non-conventional yeasts, *Y. lipolytica* is attractive for fundamental study and industrial application due to its oleaginous nature (32, 33) and robustness in extreme environments (20, 34–36). Remarkably, wildtype *Y. lipolytica* (ATCC MYA-2613) exhibited robust growth in media containing at least 10% (v/v) [EMIM][OAc] while producing 92% maximum theoretical yield of alpha-ketoglutarate from IL-pretreated cellulose (7). Currently, mechanisms of IL toxicity and superior tolerance in microorganisms are not completely elucidated (37). For instance, since prokaryotes and eukaryotes have different membrane structures, it is unclear of how their membranes are reconfigured to resist IL interference.

Given the endogenous robustness of *Y. lipolytica*, the goal of this study is to illuminate the underlying mechanisms for IL toxicity and exceptional tolerance of the wildtype strain and evolved mutant generated by ALE. Particularly, we aim to elucidate how *Y. lipolytica* restructures its membrane to resist IL disruption and modulates sterol levels to improve exceptional IL tolerance.

## Results

### Generate robust *Y. lipolytica* mutants via ALE as a basis for elucidating IL-tolerant mechanism

#### Y. lipolytica has a novel endogenous metabolism conferring exceptional IL tolerance

Since wildtype *Y. lipolytica* can grow in at least 10% (v/v) [EMIM][OAc] (7), we hypothesize that it has a novel endogenous metabolism conferring exceptional IL robustness. To test this, we performed ALE to generate *Y. lipolytica* mutants with enhanced tolerance to high concentrations of the benchmark IL, [EMIM][OAc] (Fig. 1, Step1). First, wildtype *Y. lipolytica* was grown in a medium containing 5% (v/v) [EMIM][OAc] and transferred into a medium containing progressively increased concentrations of [EMIM][OAc], 8% and 10% (v/v). Remarkably, *Y. lipolytica* was able to grow in 5%, 8%, and 10% (v/v) [EMIM][OAc] with specific growth rates of 0.063 ± 0.005 1/h, 0.056 ± 0.033 1/h and 0.060 ± 0.004 1/h, respectively, without any significant growth inhibition (Fig. 2A). When *Y. lipolytica* was transferred into a medium containing 12% (v/v) [EMIM][OAc], it initially exhibited growth inhibition with significantly reduced specific growth rate of 0.034 ± 0.001 1/h. However, after 16 generations in 12% (v/v) [EMIM][OAc], the specific growth rate was improved up to 0.080 ± 0.006 1/h and maintained for another 22 generations. Next, we increased the concentration of [EMIM][OAc] to 15% (v/v) and continued ALE. The first transfer from 12% (v/v) to 15% (v/v) reduced the specific growth rate by ~62% but cells recovered within 5 generations (0.078 ± 0.014 1/h) and the growth was maintained for an additional 23 generations. We further increased the concentration of [EMIM][OAc] to 18% (v/v) and continued serial transfers for another 106 generations. At the end of ALE (200 generations), we isolated the top performing *Y. lipolytica* mutant, YlCW001, growing in up to 18% (v/v) [EMIM][OAc] with a specific growth rate of 0.055 ± 0.006 1/h (Fig. 1, Step 2).

**Fig 1.**
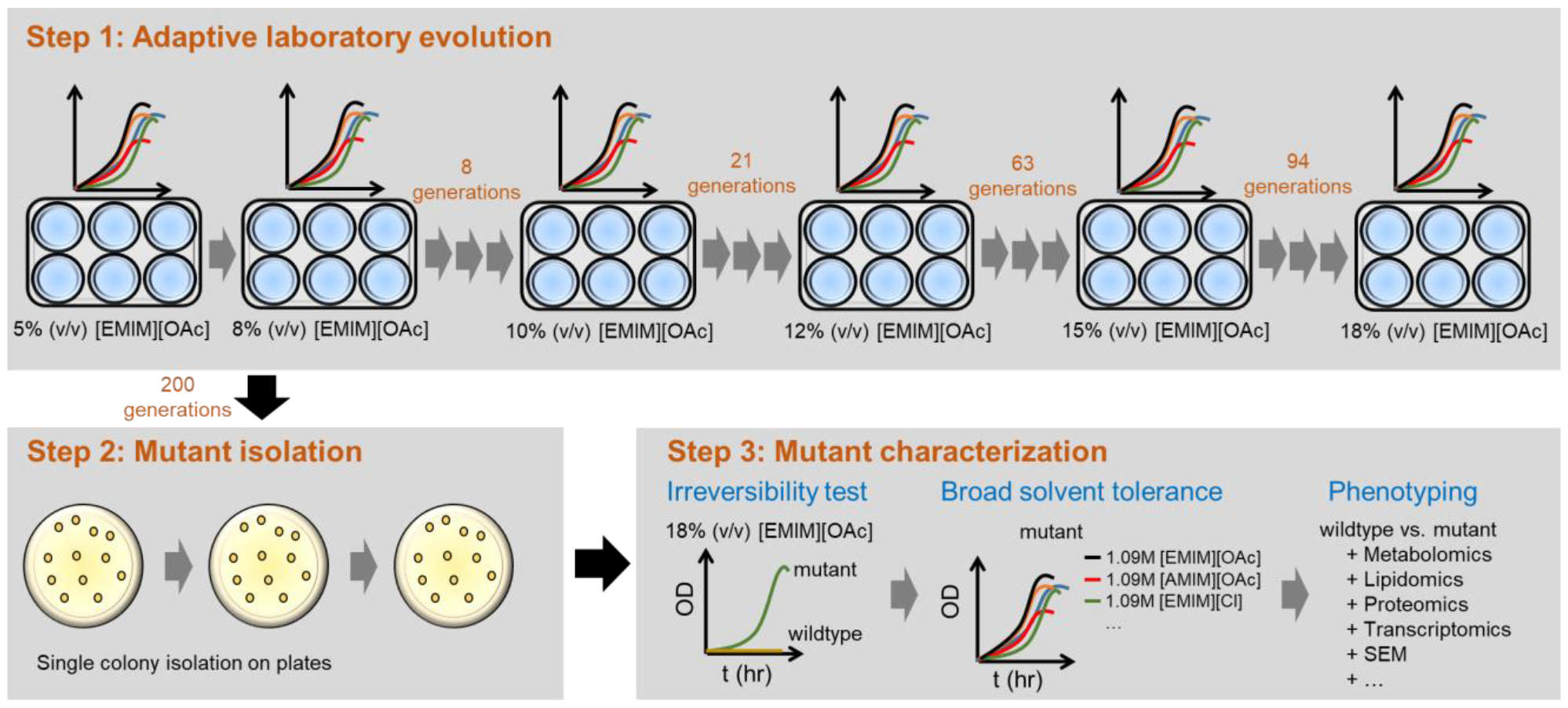
Evolutionary engineering of IL-tolerant *Y. lipolytica* strains. **Step 1**: Adaptive laboratory evolution of *Y. lipolytica* on various concentrations of [EMIM][OAc]. **Step 2**: Single mutant isolation on plates without IL. **Step 3**: Mutant characterization to elucidate the underlying mechanism of solvent tolerance. ‘

**Fig 2.**
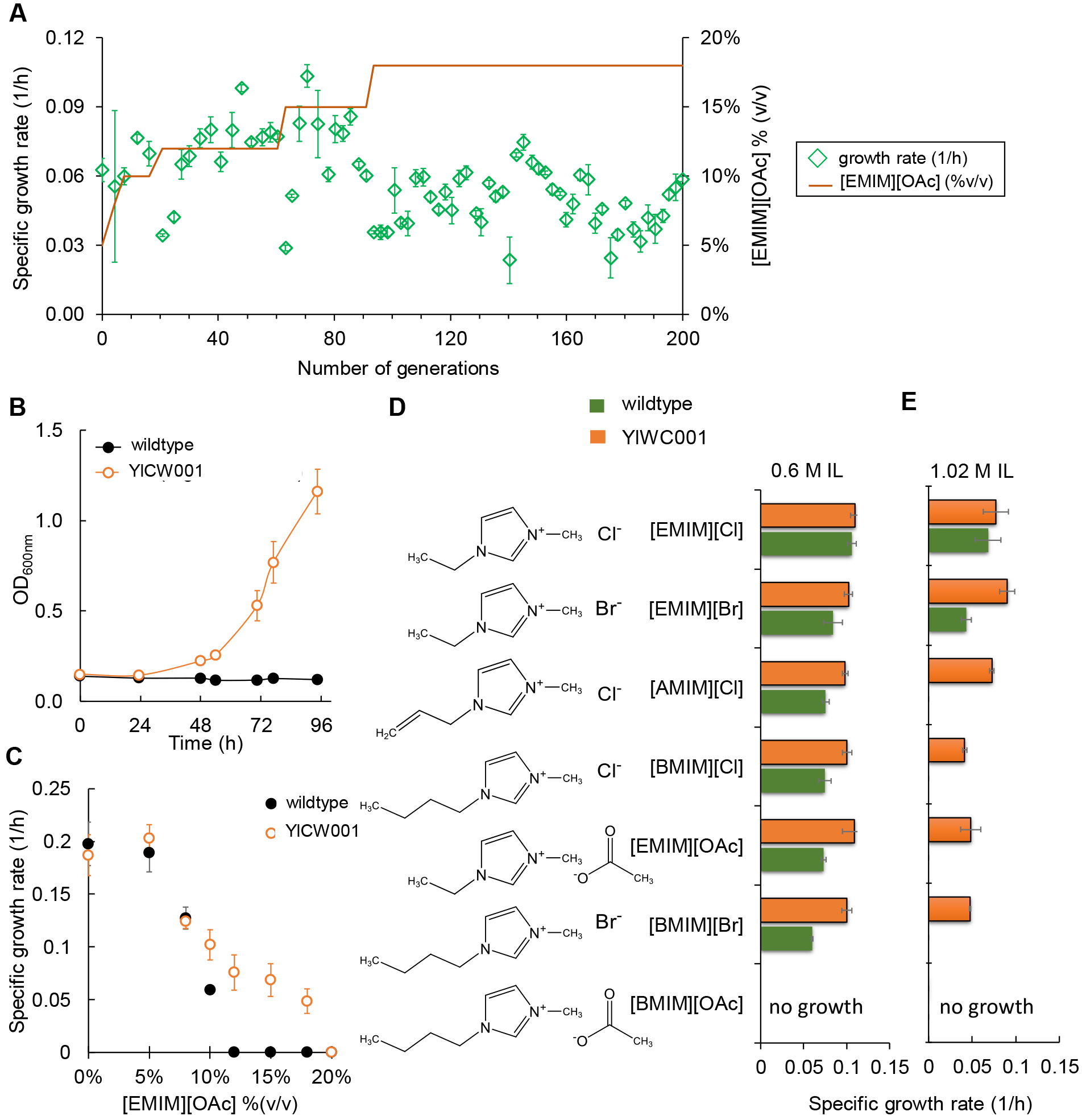
**(A)** Adaptive laboratory evolution of *Y. lipolytica* grown with increasing concentrations of [EMIM][OAc]. **(B, C)** Irreversibility testing of the evolved *Y. lipolytica* strain YlCW001 in various concentrations of [EMIM][OAc]. **(D, E)** Broad tolerance test for growth of *Y. lipolytica* wildtype and mutant at various concentrations of 0.6M and 1.09M ILs. Error bars represent the standard deviation of three independent biological replicates repeated twice (n=6). Abbreviations: [EMIM][OAc]: 1-ethyl-3-methylimidizolium acetate; [EMIM][Cl]: 1-ethyl-3-methylimidizolium chloride; [EMIM][Br]: 1-ethyl-3-methylimidizolium bromide acetate; [BMIM][OAc]: 1-butyl-3-methylimidazolium; [BMIM][Cl]: 1-butyl-3-methylimidizollium chloride; [BMIM][Br]: 1-butyl-3-methylimidazolium bromide; and [AMIM][Cl]: 1-allyl-3-methylimidizolium chloride.

#### Exceptional IL-tolerant phenotype of the evolved mutant YlCW001 is stable

To use YlCW001 for downstream characterization, we subjected it to irreversibility and stability tests (Fig. 1, Step 3). For irreversibility test, three biological replicates of wildtype and YlCW001 strains were grown in medium containing 8% (v/v) [EMIM][OAc] to lessen abrupt osmotic shock driven by IL. Mid-exponentially growing cells were then transferred into the medium containing glucose and 18% (v/v) [EMIM][OAc], and the cell growth was investigated. While growth of wildtype *Y. lipolytica* was completely inhibited, YlCW001 was able to grow with a specific growth rate of 0.059 ± 0.001 1/h (Fig. 2B), comparable to that measured from ALE in 18% (v/v) [EMIM][OAc]. These results confirmed that the improved IL-tolerance for YlCW001 is irreversible.

Further, we investigated stability of YlCW001 by reviving the frozen glycerol stock in the liquid medium containing glucose and [EMIM][OAc] (Fig. 1, Step 3). Like the irreversibility test, we tested cell growth of three biological replicates of YlCW001 in 18% (v/v) [EMIM][OAc] using glucose as a carbon source. The measured specific growth rate was similar to that from ALE and the irreversibility test (data not shown), proving that YlCW001 is a stable strain.

#### YlCW001 exhibits broad tolerance to a wide range of hydrophilic ILs

To test whether the IL-evolved strain YlCW001 exhibits broad IL-tolerant phenotypes, we investigated the following hydrophilic ILs: 1-ethyl-3-methylimidazolium acetate ([EMIM][OAc]), 1-ethyl-3-methylimidazolium chloride ([EMIM][Cl]), 1-ethyl-3-methylimidazolium bromide ([EMIM][Br]), 1-allyl-3-methylimidazolium chloride ([AMIM][Cl]), 1-butyl-3-methylimidazolium acetate ([BMIM][OAc]), 1-butyl-3-methylimidazolium chloride ([BMIM][Cl])), and 1-butyl-3-methylimidazolium bromide ([BMIM][Br]). We selected these ILs for testing because they can effectively solubilize various types of recalcitrant lignocellulosic biomass and are known to be very inhibitory to microbial growth (38, 39). Since wildtype growth was inhibited in 10% (v/v) and lethal in 18% (v/v) [EMIM][OAc] (Fig. 2C), we characterized these strains in two different concentrations of ILs, 0.6 M and 1.09 M (equivalent to 10% and 18% (v/v) of [EMIM][OAc], respectively).

Growth characterization in 0.6 M ILs shows that YlCW001 outperformed wildtype *Y. lipolytica* for all ILs except [EMIM][Cl], for which similar specific growth rates were observed (Fig. 2D). Remarkably, YlCW001 growth was not affected by all tested ILs regardless of imidazolium alkyl chain length and conjoined anion, excluding [BMIM][OAc], which at 0.6 M was lethal to both wildtype and YlCW001 (Fig. 2D). In 1.09 M ILs, wildtype *Y. lipolytica* exhibited substantial growth inhibition in [EMIM][Cl] and [EMIM][Br] and no growth in [AMIM][Cl], [BMIM][Cl], [EMIM][OAc], [BMIM][Br], and [BMIM][OAc] (Fig. 2E). Strikingly, the evolved strain YlCW001 displayed robustness in all ILs except [BMIM][OAc] confirming broad tolerance to ILs (Fig. 2E).

Inhibition of YlCW001 growth was detected in the following order: [BMIM][OAc] ≫ [BMIM][Br] ≈ [BMIM][Cl] ≈ [EMIM][OAc] > [AMIM][Cl] > [EMIM][Br] ≈ [EMIM][Cl]. While 0.6 M [BMIM][OAc] entirely inactivated growth of both wildtype and YlCW001, the mutant tolerated 0.3 M [BMIM][OAc] with a specific growth rate of 0.07 ± 0.02 1/h which remained lethal to wildtype (SI Appendix, Fig. S1).

Overall, we demonstrated that *Y. lipolytica* possesses novel endogenous metabolism conferring high IL tolerance by generating a novel, exceptionally robust *Y. lipolytica* mutant via ALE. The evolved strain YlCW001 is stable and exhibits broad tolerance towards the hydrophilic ILs in this study. This significant result provides a strong basis for elucidating the underlying mechanism of solvent toxicity in *Y. lipolytica*.

### Elucidate IL-responsive physiology and metabolism in *Y. lipolytica* strains

#### Cell membrane and morphology of the evolved mutant resist IL interference

Since ILs are known to disrupt the cell membrane (40–42), we hypothesize that the mutant YlCW001 might have adapted its membrane structure to cope with inhibitory ILs. To test this, we used scanning electron microscopy (SEM) to examine cell membranes and morphologies of both the wildtype and mutant responsive to 0% and 18% (v/v) [EMIM][OAc], 0.3M [BMIM][OAc], and 0.6M [BMIM][OAc].

As a positive control, we observed healthy morphologies for both the wildtype and mutant in no-IL media (Fig. 3A, 3B). However, when exposed to 18% (v/v) [EMIM][OAc], the wildtype developed cavities, dents, and wrinkles along the cell surface, clearly demonstrating that cell membrane and/or cell wall components were severely damaged by the IL (Fig. 3C). This phenotype is consistent with the complete growth inhibition of the wildtype observed at this high IL concentration (Fig. 2B, 2C). In contrast, the evolved strain YlCW001 displayed barely any signs of membrane damage in 18% (v/v) [EMIM][OAc] (Fig. 3D).

**Fig. 3.**
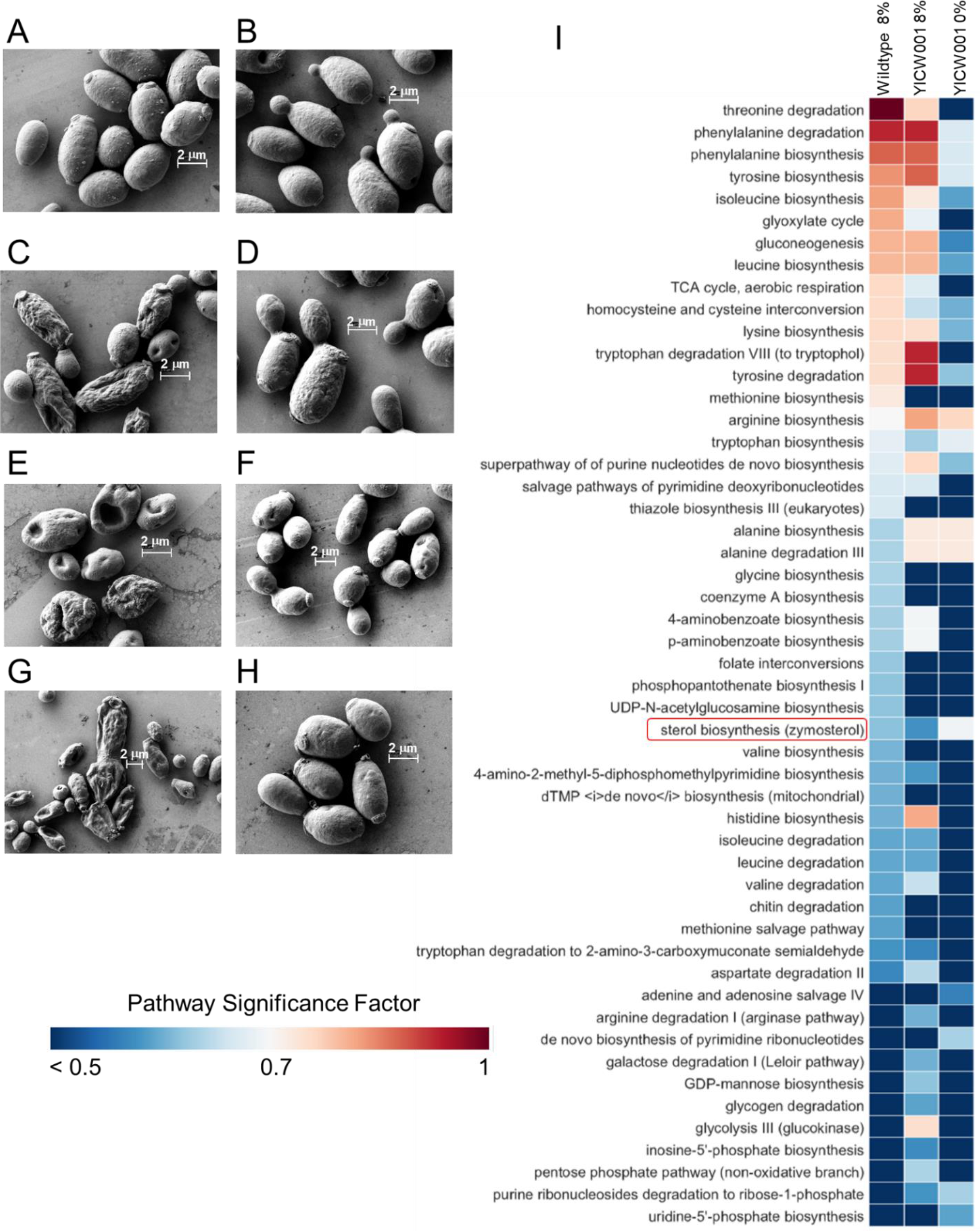
SEM imaging of the wildtype and YlCW001 cells exposed to **(A, B)** no IL, **(C, D)** 18% (v/v) [EMIM][OAc], **(E, F)** 0.3M [BMIM][OAc], and **(G, H)** 0.6M [BMIM][OAc]. **(I)** IL-responsive pathways perturbed with respect to wildtype *Y. lipolytica* growing in medium without IL with a psf score > 0.58. Biological triplicates were performed for all biological conditions and results were combined from hydrophilic metabolomics extraction (negative ionization, n = 3), and hydrophobic lipidomics extractions (negative ionization, n = 3; positive ionization, n = 3).

Likewise, when being exposed to a more toxic IL, [BMIM][OAc], the wildtype exhibited significant morphology deconstruction in both 0.3M and 0.6M [BMIM][OAc] (Fig. 3E, 3G). Strikingly, YlCW001 displayed no significant morphological defects (Fig. 3F, 3H) although growth was marginally inhibited in 0.3M and lethal in 0.6M [BMIM][OAc] (SI Appendix, Fig. S1 and Fig. 2D).

#### Intracellular metabolism of Y. lipolytica is perturbed in response to IL exposure

To demonstrate that intracellular metabolism of *Y. lipolytica* is also perturbed in IL, we performed untargeted metabolomics and lipidomics for both the wildtype and YlCW001 growing in media containing 0% and 8% (v/v) [EMIM][OAc]. Our results identified a total of 37 and 40 significantly perturbed pathways in wildtype and YlCW001, respectively, growing in 8% (v/v) [EMIM][OAc] as compared to the wildtype growing in no IL (Fig. 3I and SI Appendix, Table S1). Among these pathways, 29 were found perturbed in both wildtype and YlCW001. These pathways are mostly comprised of amino acid synthesis/degradation, nucleosides and nucleotides synthesis/degradation, but also contain vitamins synthesis (coenzyme A, thiamine, folate, and biotin biosynthesis etc.), carbohydrates biosynthesis/degradation (GDP-mannose biosynthesis, gluconeogenesis, galactose degradation, etc.), respiration (TCA cycle, glyoxylate cycle, etc.), sterols biosynthesis, and some of the central metabolic pathways responsible for generating precursor metabolites and energy (glycolysis, pentose phosphate pathway, etc.).

Furthermore, we found that YlCW001 had a total of 19 perturbed metabolic processes in media without IL in comparison to the wildtype without IL (Fig. 3I and SI Appendix, Table S1). Notably, 15 of these pathways were also perturbed in YlCW001 and wildtype growing in 8% (v/v) [EMIM][OAc] mostly enriched for amino acid biosynthesis/degradation but also included sterol biosynthesis. Overall, we identified significant perturbation of intracellular metabolism for both strains subjected to IL. While the bulk of these pathways are enriched for central carbon metabolism, our untargeted metabolomics and lipidomics identified sterols biosynthesis as the only lipid pathway significantly perturbed among all three biological conditions (wildtype 8%, YlCW001 8%, YlCW001 0%) in comparison to the wildtype without IL (Fig. 3I).

#### Remodeling of cell membrane enhanced IL-tolerance in Y. lipolytica

The outer-surface of the eukaryotic yeast is a multifaceted, permeable barrier composed of a plasma membrane (i.e., glycerophospholipids, sterols, sphingolipids) and cell wall (i.e., chitin, glucan, mannoproteins) that together, allow the cell to adapt to a variety of environmental conditions to maintain cellular homeostasis (43–46). Based on SEM images, untargeted metabolomics, and potential mechanisms of IL-toxicity (21, 47–50), we hypothesized that remodeling of cellular membrane and/or cell wall is one key IL stress-responsive process in *Y. lipolytica*. We aimed to identify critical membrane and cell wall components conferring exceptional IL-tolerance of both wildtype and YlCW001 strains by investigating IL-responsive glycerophospholipid, fatty acid, sterol, and chitin metabolism in benchmark IL, [EMIM][OAc].

#### Y. lipolytica reduced chitin in the presence of ILs

Chitin is one of the most insoluble biopolymers, even for ILs (47, 51). In *S. cerevisiae*, it has been reported that metabolism of chitin, a cell wall component known to influence membrane rigidity and elasticity, is increased upon cell wall integrity stresses (52). Since *Y. lipolytica* contains a relatively high content of chitin relative to other yeasts (e.g., ~10-15% *Y. lipolytica*; ~1-3% *S. cerevisiae*), we investigated how its membrane chitin is responsive to IL exposure. Counter-intuitively, we observed a ~2-fold reduction in chitin content for both strains upon IL-exposure (SI Appendix, Fig. S2). Of note, we were unable to detect any statistically significant differences in chitin levels between the wildtype and YlCW001 strains as they behaved similarly in 0% and 8% [EMIM][OAc]. While chitin may contribute to native *Y. lipolytica* IL-robustness, our results suggest chitin is not responsible for the enhanced IL-tolerance of YlCW001.

#### Y. lipolytica modulated glycerophospholipid composition in the presence of ILs

Upon IL exposure, the backbone and headgroups of the *glycerophospholipids* are expected to interact with both cations and anions of ILs. To understand IL toxicity and tolerance, we next investigated IL-responsive glycerophospholipid metabolism by performing targeted lipidomics on individual headgroup species (i.e., phosphatidylcholine, PC; phosphatidylinositol, PI; phosphatidic acid, PA; phosphatidyl glycerol, PG; phosphatidylserine, PS; phosphatidylethanolamine, PE; and cardiolipins, CL) for both strains in 0% and 8% (v/v) [EMIM][OAc]. In no IL media, we found that YlCW001 contained a larger amount of each glycerophospholipid species than the wildtype (Fig. 4 A-G). In 8% (v/v) [EMIM][OAc], most components of glycerophospholipids, including PC, PI, PA, PG, and CL, were upregulated in both strains, except that PE and PS exhibited different trends. The PE content of both strains remained unchanged (relative to wildtype 0%) while the PS content increased (Fig. 4E, 4F). Additionally, the mutant in media without IL contained statistically greater basal amounts of PE and PS species (p ≤ 0.01) than the wildtype.

**Fig. 4.**
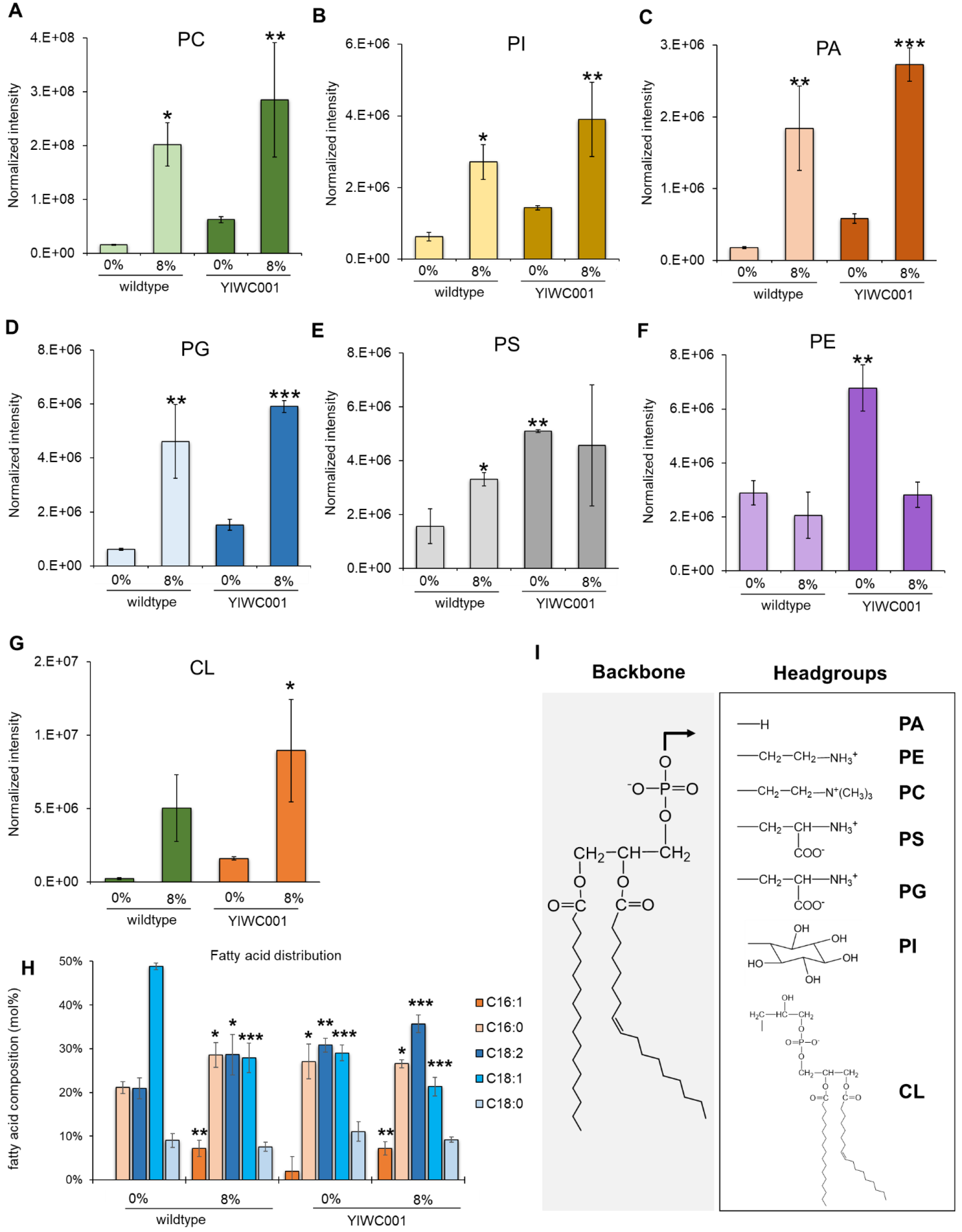
Glycerophospholipid and fatty acid reorganization in wildtype *Y. lipolytica* and YlCW001 in 0% and 8% (v/v) [EMIM][OAc]. (**A**) PC: Phosphatidylcholine (F = 11.038) (**B**) PI: phosphatidylinositol (F = 14.147); (**C**) PA: phosphatidic acid (F = 31.234); (**D**) PG: phosphatidyl glycerol (F = 29.562); (**E**) PS: phosphatidylserine (degree of freedom = 1; wildtype 8%, F = 19.953; YlCW001 0%, F = 104.891; YlCW001 8%, F = 3.097); (**F**) PE: phosphatidylethanolamine (F = 26.321); (**G**) CL: cardiolipin (F = 8.157). (**H**) Fatty acid distributions (C16:1, F = 9.501; C16:0, F = 4.768; C18:2, F = 13.4; C18:1, F = 84.261; C18:0, F = 2.839). (**I**) Chemical structures of glycerophospholipid backbone and headgroup species. All error bars represent standard deviation of biological triplicates (n=3) and statistical significance was calculated using one-way analysis of variance (ANOVA) with Holm-Sidak correction against control group, wildtype in 0% IL (degrees of freedom, 3). Symbols: “*”: *p*-value < 0.05; “**”: *p*-value < 0.01; “***”, *p*-values < 0.001.

Overall, we found IL-responsive, upregulated glycerophospholipid production of all headgroup species except PE. Interestingly, we also observed greater basal glycerophospholipid content in YlCW001 over the wildtype in 0% IL. These findings of membrane restructuring likely contribute to IL-toxicity resistance in *Y. lipolytica*.

#### ILs modulated the fatty acid composition of Y. lipolytica

Since fatty acids can modulate membrane fluidity (53, 54), we next analyzed the effect of ILs on the fatty acid profiles of wildtype and YlCW001 strains. The most striking differences were observed for C16:1 and C18:1 fatty acid moieties (Fig. 4H). We found that both strains exposed to IL induced production of C16:1 (7 mol%, *p* < 0.01) fatty acids unlike the wildtype strain without IL, which produced none. In contrast, the wildtype without IL contained mostly C18:1 fatty acids (49 mol%) while all other biological conditions shifted to the di-unsaturated C18:2 moiety (29-36 mol%, *p* < 0.02). Interestingly, YlCW001 in media without IL behaved similarly to IL-exposed strains, with significant increases in C16 and C18:2 production. We did not observe a statistically significant difference between total saturated and unsaturated fatty acid moieties in both conditions. Taken together, IL-exposed cells (and YlCW001 0%) produced C16:1 fatty acids (non-existent in wildtype 0%) with statistically significant larger ratios of C16:C18 and (C18:2):(C18:1) moieties in comparison to the wildtype without IL. Shorter chain lengths with higher degrees of unsaturation in fatty acid moieties of the cell membrane are likely expected to increase membrane fluidity (46). These results indicate that fatty acid metabolism is IL-modulated in *Y. lipolytica* and altered in YlCW001, even without IL.

#### YlCW001 increased sterols in the presence of ILs

We next investigated the functional role of sterols for IL tolerance in *Y. lipolytica* because i) the sterol biosynthesis pathway was perturbed in untargeted omics analysis, ii) sterols (e.g., cholesterol) can impede cation-insertion into the membrane (48), and iii) sterols greatly influence membrane fluidity (55, 56). In IL, we observed ~2 fold increase (*p* = 0.013) in ergosterol content of YlCW001 over the wildtype strain (Fig. 5D). Counter-intuitively, we found the largest ergosterol concentrations in both strains without IL. We were unable to identify any other sterol pathway intermediates (e.g., squalene, lanosterol, etc.), in agreement with literature concluding ergosterol as the dominant sterol in yeast (57). These results imply that IL affects sterol biosynthesis, and unlike the wildtype, YlCW001 adapted by enhancing membrane sterols in response to IL exposure.

**Fig. 5.**
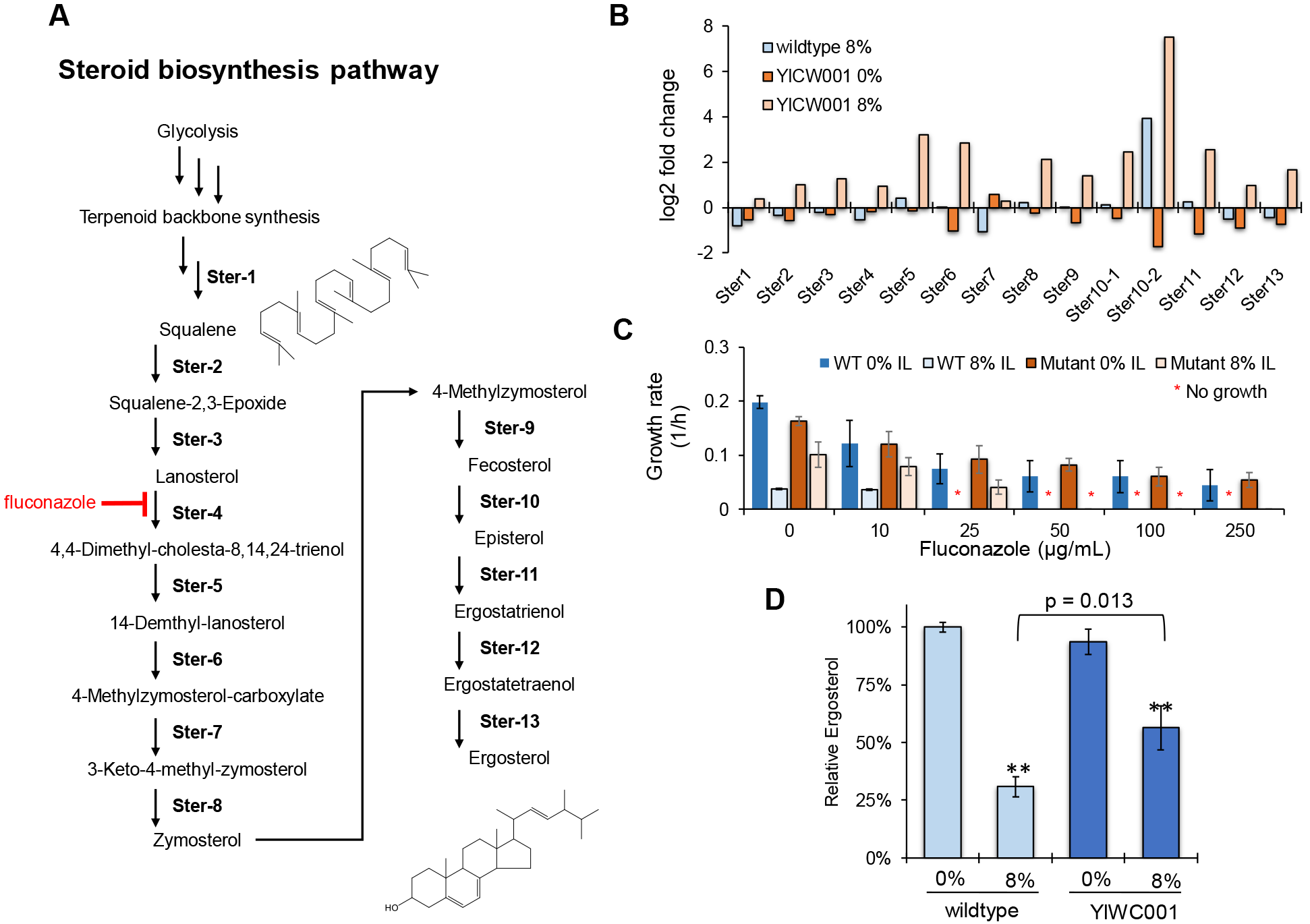
(**A**) Steroid biosynthesis pathway in *Y. lipolytica*. Fluconazole is an antifungal drug that inhibits sterol 14-demethylase (Ster4) (**B**) Differential gene expression of the steroid pathway as compared to the wildtype strain in medium without IL (Ster1, YALI0A10076g; Ster2, YALI0E15730g; Ster3, YALI0F04378g; Ster4, YALI0B05126g; Ster5, YALI0B23298g; Ster6, YALI0F11297g; Ster7, YALI0C22165g; Ster8, YALI0B17644g; Ster9, YALI0F08701g; Ster10-1, YALI0E32065g; Ster10-2, YALI0B17204g; Ster11, YALI0D20878g; Ster12, YALI0A18062g; Ster13, YALI0D19206g). (**C**) Effect of fluconazole inhibiting the steroid pathway on cell growth. Error bars represent standard deviation from three technical replicates repeated twice per biological condition (n = 6) (**D**) Relative ergosterol content of wildtype and YlCW001 strains in 0% and 8% (v/v) [EMIM][OAc]. Error bars represent standard deviation of biological triplicates (n =3) and statistical significance was calculated between the wildtype and YlCW001 in 8% IL using the Student’s t-test (t = −4.244 with 4 degrees of freedom). Abbreviation: “**”, *p*-value = 0.013.

### Sterol biosynthesis is one key IL-responsive process to improve IL-tolerance *in Y. lipolytica*

We hypothesized that ergosterol content is a critical component of the membrane contributing to the enhanced IL-tolerance of YlCW001 since we observed a greater ergosterol content in YlCW001 than the wildtype upon exposure to IL (Fig. 5D). To validate the key role of sterols, we investigated genetic and enzymatic details of how *Y. lipolytica* modulates the sterol pathway in response to IL.

We first characterized the mRNA expression levels of 14 genes in the steroid biosynthesis pathway of mid-exponentially growing wildtype and YlCW001 cells cultured in 0% and 8% [EMIM][OAc] (Fig. 5A). We found that 10 of the 14 steroid pathway genes were upregulated > 2 fold in IL-exposed YlCW001 as compared to the wildtype in 0% IL (Fig. 5B). Significantly, six of these genes, including Ster5 (YALI0B23298g), Ster6 (YALI0F11297g), Ster8 (YALI0B17644g), Ster10-1 (YALI0E32065g), Ster10-2 (YALI0B17204g), and Ster11 (YALI0D20878g), were upregulated > 4 fold in IL-exposed YlCW001 over the wildtype in 0% IL. As for the wildtype in 8% IL, we found only 1 of the 14 steroid pathway genes, Ster10-2 (YALI0B17204g), significantly upregulated against the wildtype without IL. Without IL, the steroid genes in YlCW001 were relatively constant or marginally downregulated in comparison to the wildtype.

To confirm the contribution of sterols in IL-tolerance at the enzymatic level, we next treated the wildtype and YlCW001 strains with fluconazole, a commonly used anti-fungal drug that inhibits cytochrome P450 enzyme 14α-demethylase (Ster4, Fig. 5A) critical for sterol biosynthesis (58). We characterized growth of the wildtype and YlCW001 in media containing either 0% or 8% (v/v) [EMIM][OAc] with incremental concentrations of fluconazole (Fig. 5C). We expected that fluconazole inhibits sterol biosynthesis and hence incurs the adverse effect of IL-tolerance, specifically to a greater extent in the wildtype than in YlCW001. In media containing no IL, we found that fluconazole inhibited growth for both the wildtype and YlCW001 (IC_50_ = 25 μg/mL), (Fig. 5C). In the presence of 8% (v/v) [EMIM][OAc], inhibition of fluconazole became more significant. Remarkably, YlCW001 could tolerate up to 25 μg/mL fluconazole while this concentration proved lethal for the wildtype (Fig. 5C). Taken together, the mutant in IL increased gene expression, better tolerated enzymatic inhibition of the steroid biosynthesis pathway, and elevated the end-product (i.e., ergosterol) biosynthesis in comparison to the wildtype.

## Discussion

Microbial biocatalysis in ILs is novel for biosynthesis of fuels and chemicals. For instance, the imidazolium-based ILs (e.g., [EMIM][OAc]) are effective for reducing lignocellulosic biomass recalcitrance (59) to be used for downstream fermentation but greatly inhibit microbial growth even at low concentrations (47, 60). This incompatibility presents a significant barrier for novel microbial biocatalysis in ILs. While mechanisms of IL-toxicity have been proposed (22, 29, 31), the complete picture is unclear of how cells resist to ILs and whether these cells can adapt to achieve IL-tolerance for industrial compatibility (19, 20, 27). To illuminate the mechanisms of IL toxicity and enhanced tolerance (Fig. 6), we characterized naturally IL-tolerant *Y. lipolytica* and its superior evolved mutant, YlCW001, generated by ALE (Fig. 2A).

**Fig. 6.**
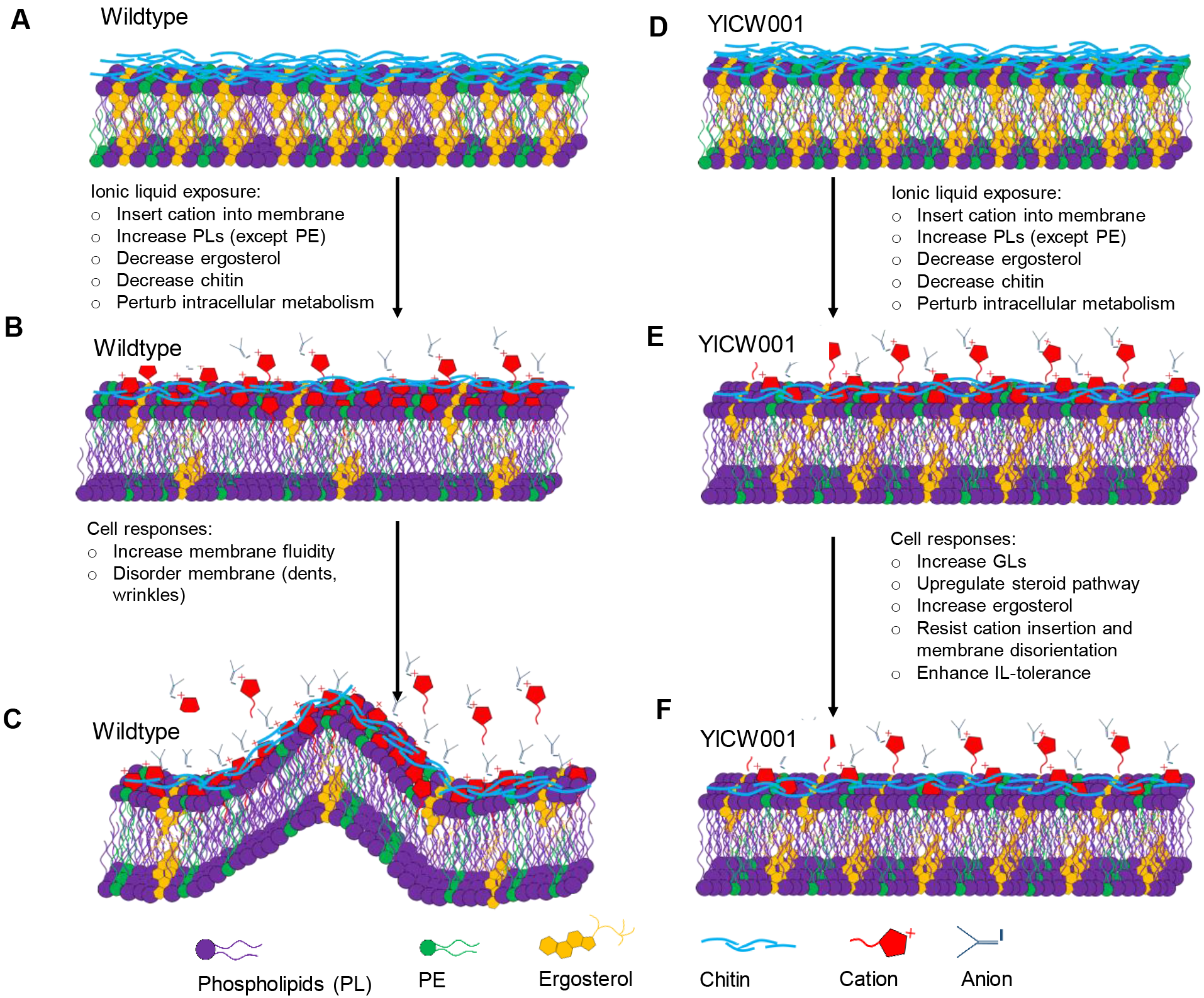
Mechanisms of IL interference and tolerance in wildtype *Y. lipolytica* and YlCW001. (**A, D**) No IL exposure. (**B, E**) IL exposure and interference. (**C, D**) Cellular membrane response.

Imidazolium-based ILs inhibit the cell by inserting their alkyl chains into the hydrophobic core of the plasma membrane (Fig. 6B, 6E, 6C, and 6F) (41, 48). ILs with longer alkyl chains become more lipophilic (21, 61) and cause greater disruption of the membrane as supported by growth rates in various ILs ([BMIM] ≫ [AMIM] ~ [EMIM]) (Fig. 2D, 2E). The conjoint anion likely re-associates with the cation imbedded into the membrane causing greater membrane disturbance (48). The tendency of an anion to interact with the cation intensifies with increasing basicity, which increases IL toxicity as demonstrated by reduced growth rates in various ILs ([OAc] ≫ [Br] ~ [Cl]) (Fig. 2D, 2E).

Consequences of IL membrane disruption increase membrane fluidity (e.g., ionic surfactant) (23, 49) and impose lateral pressure on the membrane (50, 62) as evidenced by cavities, dents, and wrinkles in SEM images (Fig. 3C, 3E, 3G) and dramatic remodeling of lipids in *Y. lipolytica* (Fig. 4). In addition, harmful interactions between the IL and membrane result in a cascade of detrimental effects on cellular processes including DNA damage, enzyme inactivation, and protein degeneration (59, 63–65), as observed by reduced sterol and chitin contents (Fig. 5D and SI Appendix, Fig. S2) and perturbation of intracellular metabolism (Fig. 3I) of IL-exposed wildtype and YlCW001 strains.

Wildtype and YlCW001 strains combatted IL toxicity in part by rewiring membrane compositions to reduce membrane permeability and bilayer buckling pressure (imposed by ILs) (66, 67). Both strains overproduced all glycerophospholipid species except PE (Fig. 4F), which is vulnerable to lateral pressure (68, 69). In contrast to the wildtype, YlCW001 is more robust because it adapts to produce more sterols, e.g., ergosterol upon IL-exposure (Fig. 5D), that function to maintain membrane fluidity and stability (55). This novel phenotype is evidenced by a significant upregulation of sterol biosynthesis genes (Fig. 5B) and improved enzymatic-tolerance to steroid-inhibiting drug, fluconazole (Fig. 5C) (58, 70). The result is also consistent with molecular simulations demonstrating sterols impede IL cations from inserting into artificial membranes (71–73).

Taken together, ILs inhibit cell growth by fluidizing the membrane and inflicting lateral pressures that destroy cellular homeostasis (Fig. 6B, 6C). Our research provides strong evidence of how intracellular processes of *Y. lipolytica* are rewired to remodel cell membranes upon IL-exposure. Comprehensive metabolic, transcriptomic, and enzymatic analyses provide strong evidence that sterols (i.e., ergosterol) are critical membrane components conferring IL-tolerance in *Y. lipolytica* and enhanced IL-robustness in YlCW001, functioning to impede cation insertion and maintain membrane homeostasis (72) (Fig. 6E, 6F). Although in this study we focused on elucidating IL-responsive metabolism specific to lipid membrane remodeling, future work will aim to determine the evolved genotype of YlCW001 to further understand IL-tolerance in *Y. lipolytica* towards the application of reverse engineering IL-robustness in diverse, industrially-relevant microorganisms.

## Materials and Methods

### Strains

*Yarrowia lipolytica* (ATCC MYA-2613), a thiamine, leucine, and uracil auxotroph, was purchased from American Type Culture Collection. The evolved strain YlCW001 was isolated after 200 generations in gradually increased concentrations of [EMIM][OAc] up to 18% (v/v).

### Medium and culturing conditions

*Growth medium*. ALE, irreversibility testing, and broad IL tolerance studies were conducted in defined media containing 6.7 g/L of yeast nitrogen base without amino acids (cat# Y0626, Sigma-Aldrich, MO, USA), 10 g/L of glucose, 100 mg/L of ampicillin, 50 mg/L of kanamycin, 30 mg/L of chloramphenicol, and various concentrations of ILs. Leucine (cat# 172130250, Acros Organics, CA, USA) and uracil (cat# 157301000, Acros Organics, CA, USA) were added to the media at concentrations of 190 mg/L and 20 mg/L, respectively.

For all other growth studies, 380 mg/L of leucine and 76 mg/L uracil were used. All ILs, including 1-ethyl-3-methylimidazolium acetate [EMIM][OAc] (>95 % purity), 1-ethyl-3-methylimidazolium chloride [EMIM][Cl] (>98% purity), 1-ethyl-3-methylimidazolium bromide [EMIM][Br] (99% purity), 1-allyl-3-methylimidazolium chloride [AMIM][Cl] (>98% purity), 1-butyl-3-methylimidazolium acetate [BMIM][OAc] (>98% purity), 1-butyl-3-methylimidazolium chloride [BMIM][Cl] (99% purity), and 1-butyl-3-methylimidazolium bromide [BMIM][Br] (99% purity), were purchased from the Ionic Liquids Technologies Inc. (IoLiTec, AL, USA). Unless specifically mentioned, all experiments were performed with biological triplicates.

*Adaptive laboratory evolution*. ALE experiment was performed by serial dilution of *Y. lipolytica* in sequentially increasing concentrations of [EMIM][OAc] in 6-well plates with 3 mL working volume using an incubating microplate shaker (cat# 02-217-757, Fisher Scientific, PA, USA) at 28°C and 350 rpm with adhesive, breathable seals to prevent cross contamination (cat# 50-550-304, Fisher Scientific, PA, USA). For each serial dilution, the top performing triplicate was transferred during mid-exponential growth phase into fresh medium at an initial optical density (OD) at 600 nm of 0.2. Increasing concentrations of [EMIM][OAc] were selected to achieve a specific growth rate ≥ 0.02 1/h. The maximum specific growth rates of all three technical replicates were calculated for each serial dilution using a minimum of three time points per replicate. After 200 generations of ALE, the top performing replicate culture was spread onto a petri plate containing defined medium with 10 g/L glucose and 20 g/L agar. The plate was incubated at 28°C for 36-48 hours. Single colonies were isolated and individually streaked onto fresh petri plates. This process was repeated once more to ensure isolation of purified colonies. Purified colonies (from three iterations of plate dilutions) were individually tested in the same growth conditions and [EMIM][OAc] concentration at which the evolved strain was originally collected to determine irreversibility. Individual cultures (from purified colonies) were collected and stored in glycerol at −80°C before streaking onto fresh petri plates, repeating three plate isolations, and retesting the irreversibility of the purified colonies to determine stability of YlCW001.

### Analytical methods

*Metabolomics*. Three biological replicates of the wildtype and YlCW001 were grown in media containing 0% or 8% (v/v) [EMIM][OAc] and collected at the late-exponential growth phase for metabolomics analysis. Samples were immediately quenched in liquid nitrogen and stored at −80°C prior to extraction. Metabolites were extracted from a minimum of 1 ×10^7^ cells in 400μL of the extraction solvent by incubating at −20°C for 20 min (74). The extraction solvent consists of 40:40:20 methanol:acetonitrile:water containing 0.1M formic acid. The soluble fraction was separated by centrifugation at 13,700 × g, 4°C for 5 minutes. To ensure complete extraction of cellular metabolites, we repeated extraction 3 times per sample. A total of 1.2 mL of solvent-soluble metabolite samples were subjected to drying under a stream of nitrogen at 4°C overnight to evaporate solvent. Lyophilized metabolites were reconstituted in 300μL of sterile water and analyzed by liquid chromatography mass spectrometer (LC-MS).

Metabolites were analyzed with an Exactive Plus orbitrap mass spectrometer (Thermo Scientific, San Jose, CA) equipped with Synergi 2.5μm Hydro-RP 100 (100 × 2.00 mm, Phenomenex, Torrance, CA, USA) set to 25°C. The LC-MS method analyzed in full scan negative ionization mode with an electrospray ionization source as previously described (75).

*Lipidomics.* Late-exponentially growing wildtype and YlCW001 cells cultured in 0% and 8% (v/v) [EMIM][OAc] were used for lipidomics study. Samples were immediately quenched in liquid nitrogen and stored at −80°C. After thawing on ice, samples were centrifuged for 3 minutes at max speed, 4°C before removing supernatant. Next, cell pellets were re-suspended in 800uL of 0.1N hydrochloric acid:methanol 1:1 with 400uL of chloroform and disrupted with glass beads using a mini bead beater for 5 minute intervals until > 95% of cells were visually disrupted. Disrupted cells were vortexed and centrifuged at 4°C for 5 minutes before extracting the organic phase into glass vials. Finally, samples were dried under a stream of nitrogen overnight at 4°C and reconstituted in 300uL of 9:1 methanol:chloroform before transferring into auto-sampler vials. Lipid extracts were analyzed in positive and negative ionization modes with an Exactive Plus orbitrap mass spectrometer (Thermo Fisher Scientific, San Jose, CA) equipped with an electrospray ionization probe and a Kinetex HILIC column (150 mm × 2.1 mm, 2.5 μm) (Phenomenex, Torrance, CA, USA) as previously described (76) except that lipid features were verified with external standards instead of fragments.

*Untargeted LCMS analysis*. Metabolomics and lipidomics raw files created by Xcalibur were imported into XCMS online and analyzed in pairwise jobs against the control data set, wildtype in 0% IL (77). XCMS resulting metabolic features were exported and features with intensity fold changes < 2 were removed. These pairwise sets of ‘perturbed’ metabolic features were analyzed with metaboanalyst (78) using ‘MS peaks to pathways’ tool with significant feature *P*-value cutoff = 0.05. This tool uses mummichog which algorithmically utilizes known metabolic pathways and networks to predict metabolites and pathways without prior identification of metabolites (79). The resulting pathway files from ‘perturbed’ pairwise feature sets were exported and pathways with less than two significant features (i.e., metabolite features with *P*-values < 0.05) were removed. Next, we filtered the pathways by defining a new parameter, pathway significance factor (psf) (equation 1), to account for i) total number of metabolites in the pathway (p_size_), ii) total identified metabolite features (p_features_), and iii) total number of significant metabolite features (p_significant_) identified for each pathway.

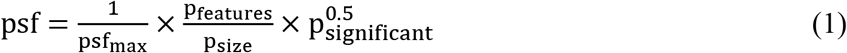

In our analysis, we chose a psf cutoff value of 0.58 to illustrate the top 15% most significantly perturbed pathways identified from our untargeted LCMS analysis.

*Targeted LCMS analysis*. Lipidomics raw data files created by Xcalibur were converted to open source mzML format using the ProteoWizard software (80). MAVEN software (Princeton University) was applied to performed retention time correction for each sample and used to manually select known lipids based on retention time and mass (81, 82). Glycerophospholipid headgroup species were analyzed individually and extracted signal intensities were corrected by cell optical density. The corrected intensities for each head group class were summed to visualize changes for each glycerophospholipid on a macro scale.

*Fatty acid quantification*. Three biological replicates of wildtype and YlCW001 cells were grown in 0% and 8% (v/v) [EMIM][OAc] until mid-late exponential phase and cell pellets were stored at −20°C. The equivalent cell mass of 2 OD_600nm_ was washed once with 0.05 M sodium phosphate solution and incubated in 2:1 (v/v) chloroform:methanol solution overnight at 4°C. 200 μL of chloroform was extracted and mild methanolysis was performed as previously described (83). Briefly, 1.5 mL of methanol, 0.3 mL of 8% methanolic HCL solution and 50 μL of 2 mg/mL pentadecanoic acid was added to ensure complete transesterification and incubated overnight at 55°C. After cooling to room temperature, 1mL of hexane containing 0.005 mg/mL pentadecanoic acid ethyl ester as internal standard and 1mL of water was added prior to extracting 250 μL of hexane for GCMS detection of fatty acid methyl esters.

*Sterol quantification*. Three biological replicates of wildtype and YlCW001 cells were grown in 0% and 8% (v/v) [EMIM][OAc] until mid-late exponential phase and cell pellets were stored at −20°C. Sterol quantification was performed with the equivalent cell mass of 5 OD_600nm_ as previously described (84) using a HP-5MS 30 m × 0.25 mm i.d. × 0.25 μm column (Agilent Technologies, USA) for separation of sterols.

*Chitin determination*. The relative quantity of chitin was analyzed for the wildtype and YlCW001 late-exponentially growing cells in medium containing 0%, 2%, 5%, 8% and 10% (v/v) [EMIM][OAc]. Cell pellets were washed twice with water before resuspending 1 OD_600nm_ of cell mass in 1 mL of water containing 50 μg/mL calcofluor white (CFW) (cat #18909, Sigma-Aldrich), which binds specifically to chitin (85, 86). Samples were incubated at room temperature for 15 minutes at 650 rpms in a thermomixer (cat #5382000023, Eppendorf). Next, stained cells were washed twice with water to remove excess CFW prior to fluorescence (excitation: 360/40nm, emission: 460/40nm) and absorbance (OD_600nm_) measurements. Results were calculated by normalizing fluorescence intensity by respective sample OD values. Chitin determination experiments were conducted at least twice for each biological condition in technical replicates per experiment (wildtype 0%, n = 18; wildtype 2%, n = 6; wildtype 5%, n = 15; wildtype 8%, n = 12; YlCW001 0%, n = 25; YlCW001 2%, n = 6; YlCW001 5%, n = 15; YlCW001 8%, n = 29; YlCW001 10%, n = 6).

*Scanning electron microscope (SEM)*. Wildtype and YlCW001 cells were inoculated at 1 OD_600nm_ in 0%, and 18% [EMIM][OAc] and 0.3M and 0.6M [BMIM][OAc] in 6-well microtiter plates using an incubating microplate shaker at 28°C and 350 rpm. After 24 hours, cells were collected, immediately washed once with water and incubated in 2% glutaraldehyde containing 10 mM sodium phosphate buffer overnight at 4°C. Fixed samples were washed 3 times with water before post-fixing in 2% osmium tetroxide for 1 hour at room temperature. The cell pellets were placed on silicon chips and dehydrated with successive ethanol baths (50%, 60%, 70%, 80%, 90%, 100%) for 15 minutes each. Finally, the dehydrated samples were dried using critical point drying in carbon dioxide at 1100 psi and 32°C before SEM imaging with Zeiss Argula using SEM2 detector.

*RNA-sequencing of steroid biosynthesis pathway*. In order to investigate response of sterol biosynthesis genes in response to IL, we quantified expression level of the 14 steroid genes annotated in the KEGG database (87) using RNA-sequencing. Three biological replicates of wildtype and YlCW001 were harvested at the mid-exponential growth phase from 0% and 8% (v/v) [EMIM][OAc] and immediately quenched in liquid nitrogen before storing samples at −80°C. Total RNA was purified using a Qiagen RNeasy mini kit (Cat no. 74104, Qiagen Inc, CA, USA). Filtered RNA-sequencing reads were imported and analyzed within the CLC genomics workbench version 11.0.1 (https://www.qiagenbioinformatics.com/) which was also used to calculate differential expression of steroid biosynthesis genes against the wildtype in 0% IL.

*Fluconazole treatment to investigate importance of sterol biosynthesis for enhanced IL tolerance*. To investigate correlation of sterol biosynthesis with enhancement of IL tolerance, we compared growth of the wildtype and YlCW001 in media containing 0% or 8% (v/v) [EMIM][OAc] and various concentrations of fluconazole (cat# TCF0677, VWR) which inhibits fungal cytochrome P450 enzyme 14α-demethylase (Ster4, Fig. 5A) required for sterol biosynthesis (58). First, we cultured the wildtype and YlCW001 in media without IL until the mid-exponential phase. Then, cells were washed by water and resuspended in the fresh media containing 0, 10, 25, 50, 100, and 250 μg/mL fluconazole. Finally, the same conditions were tested with addition of 8% (v/v) [EMIM][OAc] to observe the requirement of sterol incorporation into the plasma membrane for IL tolerance. Results were obtained by calculating maximum specific growth rates for each biological condition. Sterol validation by fluconazole treatment experiments were conducted twice for each biological condition with sacrificial, technical replicates using 96-well microtiter plates and Duetz sandwich cover (cat # SMCR1296, Kuhner) incubated at 28°C and 400 rpms.

*Statistics*. Statistical analysis was performed with SigmaPlot v. 14 software using one-way analysis of variance (ANOVA) with Holm-Sidak correction or Student’s t-test between biologically relevant conditions where noted.

## Acknowledgement

This research is funded by the National Science Foundation (NSF CBET#1511881). We would like to thank Dr. Stephen Dearth (Biological and Small Molecule Mass Spectrometry Core, UTK) for assistance in metabolomics and lipidomics experiments, Dr. John Dunlap (Advanced Microscopy Imaging Center, UTK) for assistance in SEM experiments, and Mr. Piet Jones (Bredesen Center, UTK) for technical support. The RNAseq work conducted by the U.S. Department of Energy Joint Genome Institute, a DOE Office of Science User Facility, is supported by the Office of Science of the U.S. Department of Energy under Contract No. DE-AC02-05CH11231.

